# MgO nanoparticles coated with Polyethylene Glycol as carrier for 2-Methoxyestradiol anticancer drug

**DOI:** 10.1101/588939

**Authors:** Aline Alfaro, Andrea León, Eduardo Guajardo, Patricia Reuquen, Francisco Torres, Mario Mery, Rodrigo Segura, Paula A. Zapata, Pedro A. Orihuela

## Abstract

Novel Magnesium Oxide (MgO) nanoparticles (NPs) modified with the polymer poliethylene glycol (PEG) were synthesized as carrier for the anticancer drug 2-Methoxyestradiol (2ME) to improve its clinical application. The functionalized NPs were characterized by Infrared spectroscopy with Fourier transform to elucidate the vibration modes of this conjugate, indicating the formation of the MgO-PEG-2ME nanocomposite. The studies of absorption and liberation determined that MgO-PEG-2ME NPs incorporated 98.51 % of 2ME while liberation of 2ME was constant during 7 days at pH 2, 5 and 7.35. Finally, the MgO-PEG-2ME NPs decreased the viability of the prostate cancer cell line LNCap suggesting that this nanocomposite is suitable as a drug delivery system for anticancer prostate therapy.

## Introduction

Cancer is one of the diseases with higher prevalence in the population, being the second cause of death in 2015, with a total of 8.8 million globally. It is expected that due to the growth and aging of the population in the next two decades 22 million of people will be diagnosed annually with this pathology [1, 2]. At global level the most common kinds of cancer are; lung, breast, duodenum and prostate [2, 3]. Prostate cancer is the most common type of carcinoma in male from developed and developing countries [4]. The current therapies for prostate cancer are focused in surgery, radiation and hormonal treatment. Unfortunately, survival prognostic for patients is very poor because many patients will suffer from recurrence and subsequent metastasis [4–6]. Therefore, it is necessary development new drugs and therapies that will be effective for the treatment of prostate cancer.

2-Metoxyestradiol (2ME) has antitumor activity in several types of cancer of the reproductive tract as prostate, cervix, ovary or endometrium. 2ME exerts its anticancer activity via anti-proliferative, apoptotic or antiangiogenic effects on tumor cells [7]. Despite to be considerate as a promising anticancer drug it has an unfavorable kinetic with a low solubility in water; Thus, it is necessary to find new ways to facilitate its administration to the human body. In this context, the nanoparticles (NPs) as drug carriers can play a fundamental role to improvement biological parameters. Actually, it has been proposed that polymeric NPs [8] or TiO_2_ NPs coated with polyethylene glycol (PEG) could be useful tools to load 2ME [9]. In the searching for new NPs suitable for medical use, MgO NPs are also an excellent candidate because they are bio-friendly [10]. It has been shown that MgO NPs are not toxic for a variety of human cell lines at concentrations under 300 µg/ml [10]. MgO NPs has been also used as a carrier for the antincancer drug doxorubicin indicating its utility for a controlled system of drugs delivery [11, 12]. In this work, we first performed the physicochemical characterization of MgO NPs coated with PEG and loaded with 2ME (MgO-PEG-2ME). The efficiency of absortion and liberation of 2ME was then analyzed. Finally, the effect of MgO-PEG-2ME NPs on the prostate cell line LnCap was assessed.

## Materials and Methods

### Synthesis MgO nanoparticles

The MgO NPs were obtained by the sol-gel method route assisted with cetyltrimethyl ammonium bromide C_19_H_4_2BrN (CTAB) as a surfactant to reduce the agglomeration of the NPs [13]. 1:1 molar solution of magnesium acetate, Mg(CH_3_COO)_2_ 4H_2_O (99,5 %, MERK, USA) and tartaric acid C_4_H_6_O_6_ to (99,5 %, MERK, USA) was prepared in ethanol and added dropwise over 10 ml of a 0.001 M of CTAB in water at 60°C. The solution was stirred vigorously for 20 hours to achieve gel formation. Once the gel is formed, it was dried and before calcined at 600°C for 6 hours to give MgO [14].

### MgO nanoparticles functionalized with PEG and 2ME loading

MgO NPs were covered with poliethylene glycol 600 (PEG600; Sigma Aldrich) using the agitation method. For this, 50 mg of MgO NPs were dispersed on 50 ml MiliQ water and stirred for 1 hour, then 88 *µ*l of PEG 2mg/ml was added and stirred by 2 hours and centrifuged for 30 minutes at 4500 *g*. The supernatant was discarded and the solid phase was dried on a heat plate (Memmert, Germany) at 60°C. Then, 2ME 1 mg was added to 1 mg of MgO-PEG NPs and stirred for 2 hours and centrifuged at 10.621 *g* for 1 hour at 10°C. Then, the solid phase was rinsed and dried on a heater plate (Memmert, Germany) at 60°C.

### Characterization techniques

The morphology and size of the NPs were determined by Transmission Electron Microscopy (TEM). The NPs were supported in a copper mesh covered with carbon (Support Films, Carbon Type-B, Ted Pella, inc.). The observations were performed with a TEM HT7700 (Hitashi, Japan) at an acceleration voltage of 80 kV. The mean diameter particle of the NPs was obtained measuring at least 120 particles using the ImageJ software (National Institute of Health, USA). The functionalization of MgO-NPs by PEG and conjugation with 2ME were examined by Attenuated Total Reflectance Infrared Fourier-transform (FTIR) spectroscopy (ATR-FTIR). The FTIR spectra was collected in the 4000-1000 cm^-1^ range, with a resolution of 4 cm^-1^ at room temperature by using a Thermo Nicolet IS10 spectrometer provided with single bounce Ge crystal Smart-iTR accessory.

### Zeta potential

The zeta potential of the NPs were analyzed by dynamic light scattering in the Zetasizer Nano ZS DST1070 cell (Malvern Instruments, UK) [15]. Preparations were dissolved in 1 ml phosphate buffered saline at pH 7.35. The Samples were evaluated in triplicate.

### Ultra-high performance liquid chromatography (UPLC)

UPLC was performed using an Acquity system (Waters, Milford, MA, USA) equipped with a binary solvent delivery pump, an autosampler and a tunable UV detector. Chromatographic separation was performed using a Waters Acquity BEH C18 column (50 × 2.1 mm, 1.7 mm). The mobile phase was a 70:30 (v:v) mixture of methanol and water at a flow rate of 0.4 ml/min. Detection was performed at a wavelength of 290 nm using a 10 µl injection volume; the mobile phase of water and methanol was maintained at 27°C. The internal chromatographic standard solutions (1, 5, 10, 50 and 100 mg/ml) were freshly prepared in a volumetric flask along with the mobile phase [16].

### Efficiency of absorption and liberation

The 2ME entrapment efficiency was analyzed using an extraction method described in our previously word [9]. 1 mg of MgO-PEG-2ME NPs was dispersed in 1 ml PBS, and agitation on an orbital shaker at 100 rpm. Samples were taken at 5 min, 30 min, 1 h, 3 and 6 hours at 37°C. The 2ME concentration was determined by UPLC using a calibration curve. To measure 2ME release, 1 mg of MgO-PEG-2ME NPs underwent rapid equilibrium dialysis (Thermo Scientific, see manufacturers instructions) through sequential bag dialysis at 37°C with gentle shaking in 15 ml of PBS (pH 2, 5 and 7.35). At each sampling time, 1 ml of the supernatant was removed and replaced with an equivalent volume of PBS. The supernatants were analyzed by UPLC to determine 2ME release.

### Cell Culture

LNCap cells were grown in DMEM medium (Hyclone, USA) supplemented with sodium pyruvate 1 mM, 10% heat-inactivated fetal bovine serum, 100 UI/ml penicillin, 100 *µ*g/ml streptomycin under 5% CO_2_ in 95% of air in a cell culture incubator at 37°C. The cells were used when reach to 70-80 % of confluency. For all experiments, 2.5 × 10^3^ cells/well were seeded.

### Treatments and Measurement of cell viability

LNCaP cells were treated with nanoparticles of MgO, MgO-PEG or MgO-PEG-2ME at a concentration equivalent to 5 *µ*M of 2ME. The cell viability was assessed by the viability assays 3-(4,5-dimetiltiazol-2-il)-5-(3-carboximetoxifenil)-2-(4-sulfofenil)-2H-tetrazolio (MTS) using the Cell-Titer 96^*c*^ AQueous Non-Radioactive Cell Proliferation Assay kit (Promega, Madison, USA) according to manufacturer’s instructions. LNCaP cells were grown on 96-well assay plates and 24, 48 or 72 hours post-treatment, 20 *µ*l of MTS reagent provided by the kit was added to each well. After incubation, the absorbance value at 490 nm was obtained using an ELISA plate reader (Tecan Group Ltd. Mnnedorf, Switzerland). As positive control of cytotoxicity we added hydrogen peroxide (H_2_O_2_) 0.08% dissolved in 4 *µ*l of culture medium. Furthermore, as control we used a solution of 2ME 5 *µ*M. Ethanol 0.01 % was used as vehicle of the nanoparticles and 2ME.

### Statistical analyses

Cellular viability assays were performed in triplicate. The data were analyzed using GraphPad Prism (GraphPad Software, San Diego, CA, USA). When correspond, all data are presented as mean standard error. Overall analysis was done by Kruskal-Wallis test followed by Mann-Whitney U test for pair-wise comparisons when overall significance was detected. Differences were considered significant at P < 0.05.

## Results and Discussions

### Characterization of MgO, MgO-PEG and MgO-PEG-2ME NPs

Fig 1A and 2 reveals that the MgO NPs are lightly agglomerated forming small particles in a narrow size distribution with a mean value of 15.7 ± 4.3 nm. When PEG and 2ME were add to MgO NPs, the size mean value increase with the size around 16.0 ± 10.6 nm (Fig.1B and 2) and 93.31 ± 74.0 nm (Fig1.C and 2) for MgO-PEG and MgO-PEG-2ME respectively. The histogram distribution for pure MgO-NPs, MgO-PEG and MgO-PEG-2ME is shown in the Fig 2.

**Fig 1.**
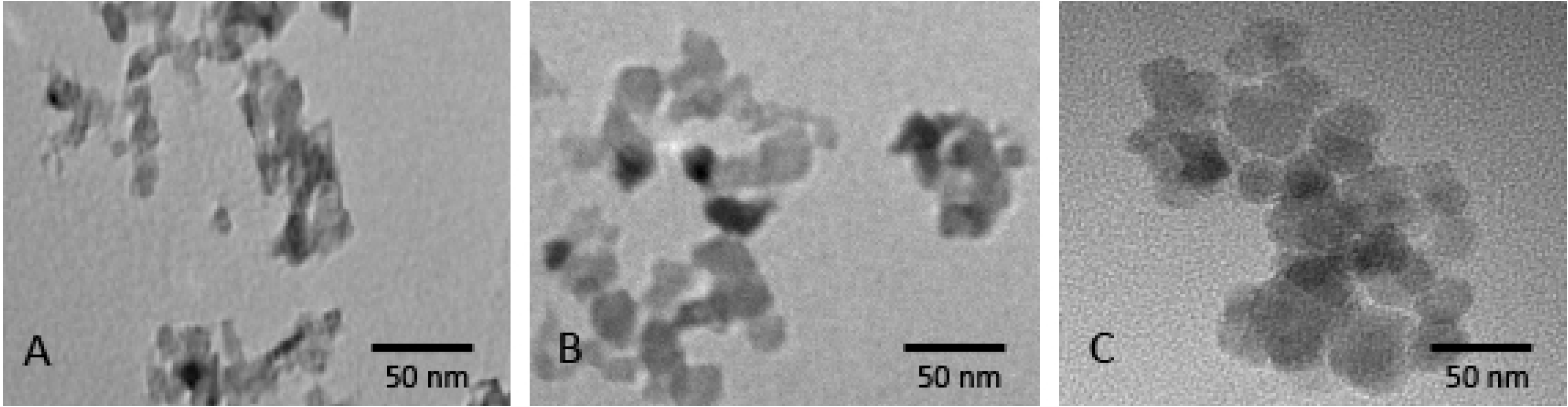
Nanoparticles Photomicrographs. Representative nanoparticles photomicrographs: (A) MgO, (B) MgO-PEG and (C) MgO-PEG-2ME. Note that MgO-PEG-2ME are bigger than other nanoparticles.

**Fig 2.**
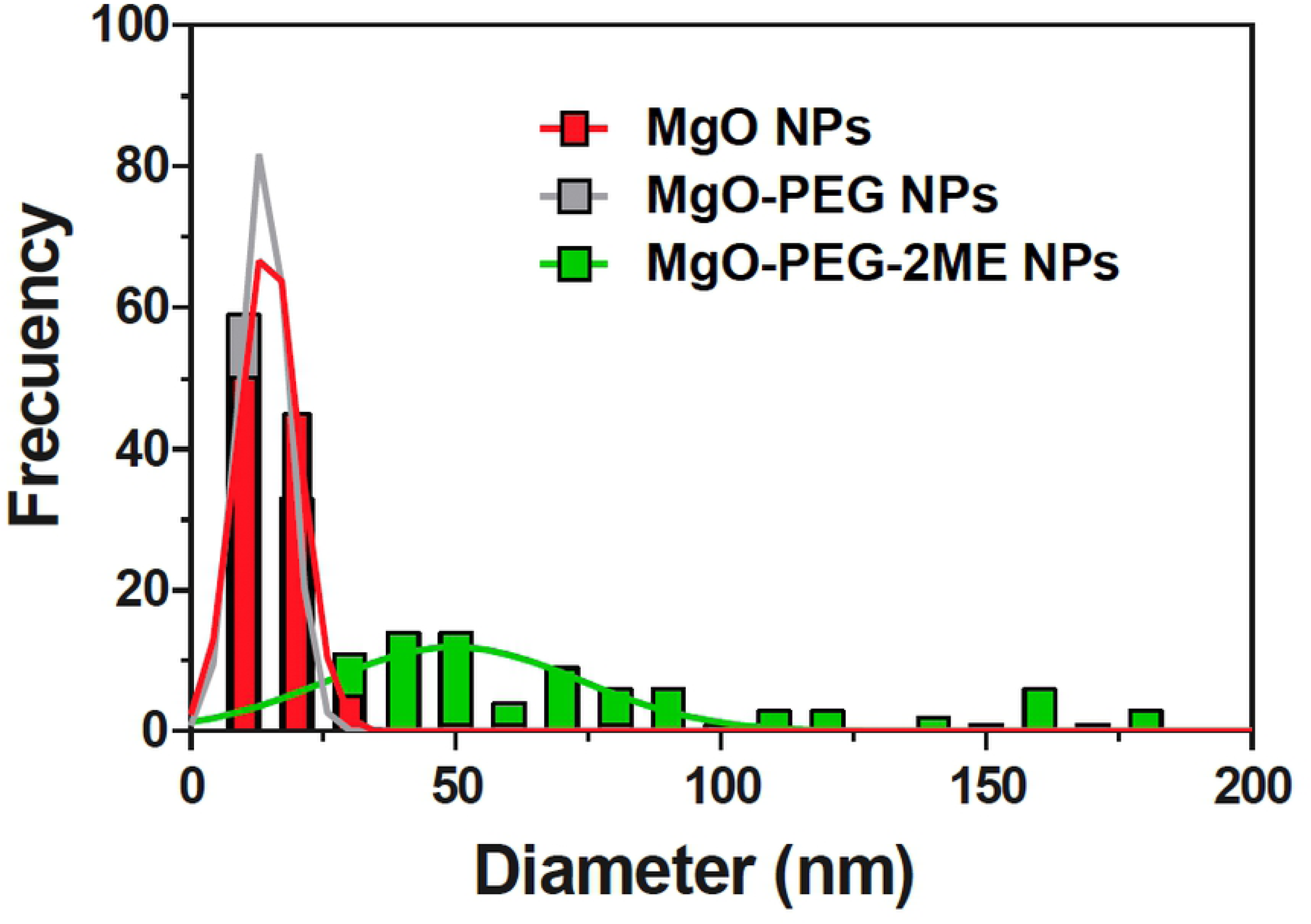
Histogram. Histogram distribution of the MgO, MgO-PEG and MgO-PEG-2ME NPs.

The size of the NPs is an important parameter that determines his biocompatibility; our results for TEM analysis show a wide size distribution, which could be optimal for the internalization into the intracellular space via different mechanisms as endocytosis, phagocytosis and pinocytosis. Particularly, endocytosis mediated by clathrin and caveolae induce a greater accumulation of NPs inside the cells [17]. In addition, it must be considered that for each NPs that is able to be transported across the cell membrane exist an ideal radius that allows a rapid internalization. As this radius is approximately 90 nm for spherical conjugate NPs, we can assume that our NPs are suitable for cell internalization [18]. Other variables that determine the biocompatibility of the NPs is the superficial charge due to chemistry modification. The results of the zeta potential measurements for MgO, MgO-PEG and MgO-PEG-2ME NPs are shown in the table 1. We can observe a negative zeta potential increasing from −30 meV to −28.3 and −22.0 when PEG and 2ME are added suggesting that these NPs could be useful for biological applications. In this context, It has been shown that NPs with positive zeta potential could be more deleterious because induce platelet aggregation that can cause thrombosis or they can interact with membrane phospholipid or proteins disrupting stability of the cell surface [18].

**Table 1.** Mean *±* Zeta Potential.

| NPs | Zeta Potential (mV) |
| --- | --- |
| MgO | $-30.7 \pm 0.6$ |
| MgO-PEG | $-28.3 \pm 3.5$ |
| MgO-PEG-2ME | $-22.0 \pm 10.8$ |

Mean ± Zeta Potential for Nanoparticles of MgO, MgO-PEG and MgO-PEG-2ME NPs.

To characterize and determine functional groups and modifications, FTIR spectroscopy was performed on pure MgO-NPs, PEG and 2ME as well as in MgO-PEG, MgO-PEG-2ME NPs and PEG-2ME. Fig 3A shows the spectrum of the MgO, PEG, MgO-PEG, MgO-PEG-2ME. For MgO, three principal bands are found; at 3700 cm^-1^ belong to the O-H group, at 1400 cm^-1^ corresponding to Mg-O streching vibration [19, 20] and at 846 cm^-1^ which has been attributed to the formation of cubic phase of MgO [19]. The FTIR PEG shows the characteristic bands at 2929 cm^-1^, 2888 cm^-1^, 1342 cm^-1^, 1242 cm^-1^ and 1100 cm^-1^ that have been described by [21, 22]. When PEG is added to NPs we can observe that the bands are modified, with respect to pure MgO-NPs mainly at 1098 cm^-1^ (C-O streching vibration) [19], this band is shifted at 1081 cm^-1^ from their original position in pure PEG, exhibiting hydrogen-bonding nature and suggests that PEG interaction with the surface of MgO-NPs [19]. The FTIR spectrum of pure 2ME exhibit characteristic bands occurring at 3417 cm^-1^, 3182 cm^-1^, 3000 cm^-1^, 2963 cm^-1^, 2907 cm^-1^, 2809 cm^-1^, and 1600 cm^-1^, and also in the ranges between 1500-1400 cm^-1^ and 1300-1000 cm^-1^, the last bands are the fingerprint of 2ME, that have been described previously by our research group [9].

**Fig 3.**
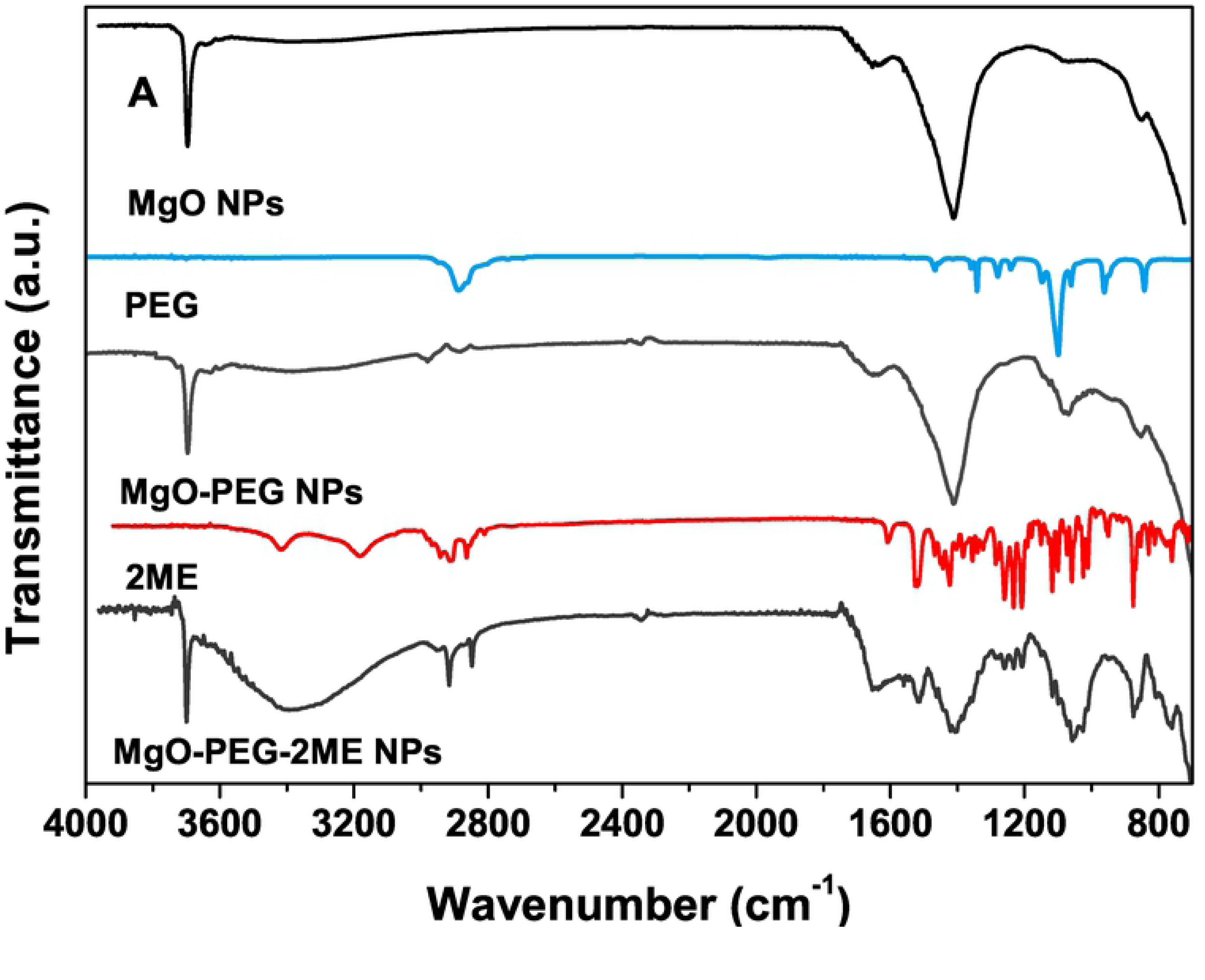
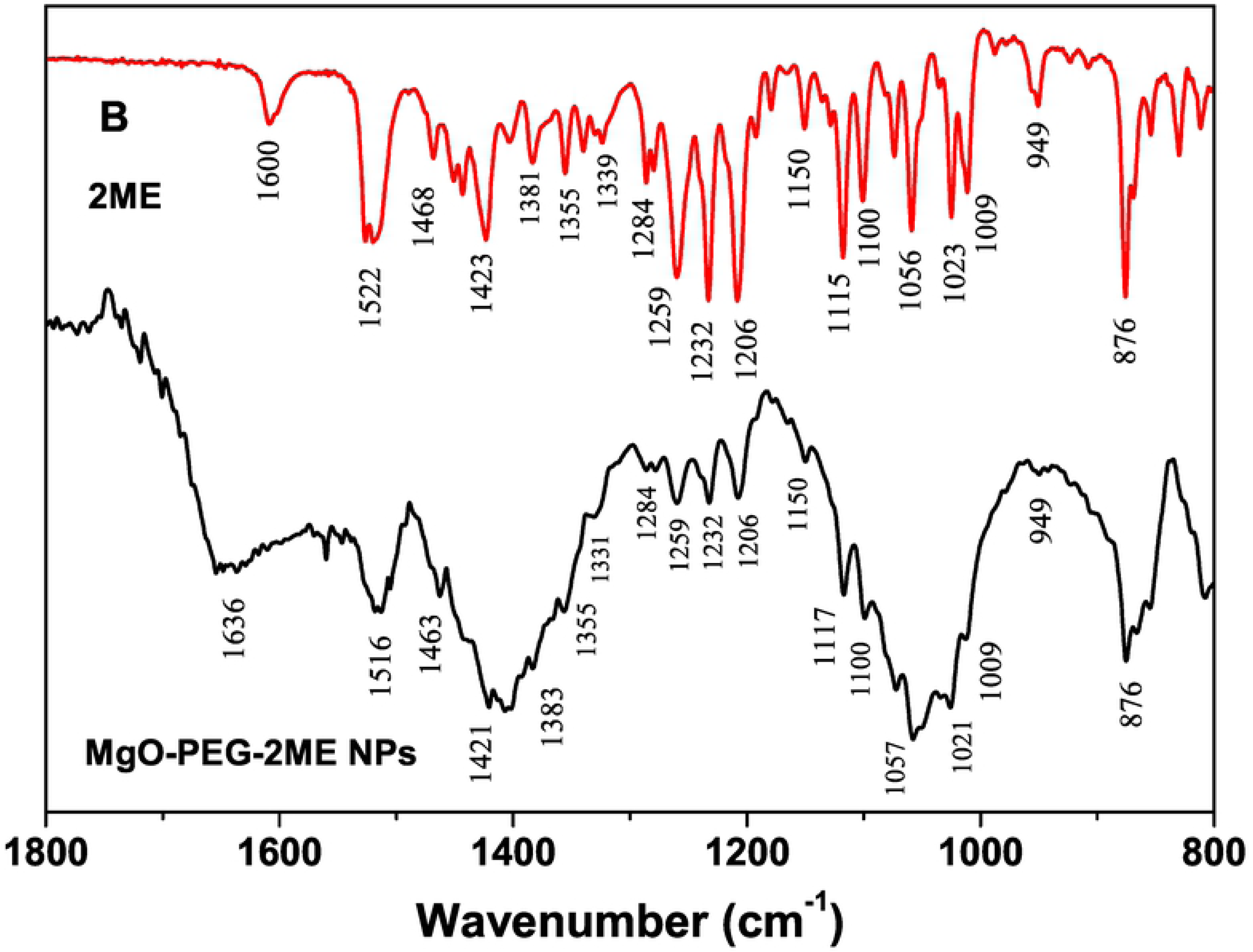
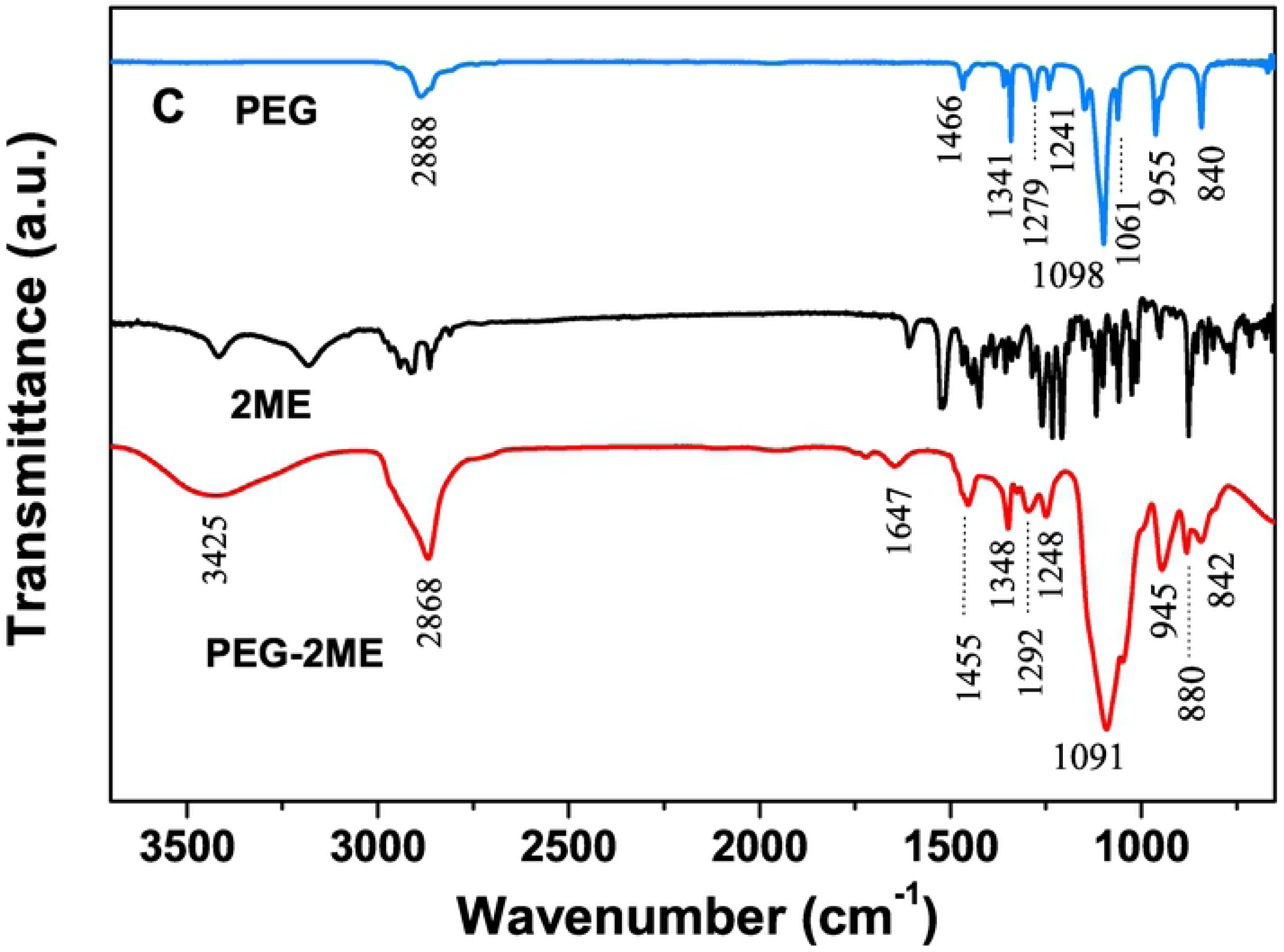
Fourier-Transform Infrared (FTIR) Spectra. (A) Fourier-transform infrared (FTIR) spectra for: MgO, PEG, MgO-PEG, 2ME, and MgO-PEG-2ME. (B) 2ME and MgO-PEG-2ME NPs spectra in the range of [1800-600 cm^-1^]. In this zone we can distinguish the principal functional groups of 2ME and the new bands that appear in MgO-PEG when is conjugate with 2ME, where the mains bands are labeled. (C) PEG, 2ME and PEG-2ME; this figure shows the modified band of PEG when is conjugate with 2ME, where the mains bands are labeled.

In the FTIR spectrum of MgO-PEG-2ME NPs, we found several changes including new bands at 2861 cm^-1^ and 2919 cm^-1^ belonging the C-H stretching vibration in CH, -CH_2_, -CH_3_. The band at 3382 cm^-1^ is related with the stretching vibration of hydroxyl group, this band is broadened with an increased intensity with respect to pure 2ME. The band localized between 1500-1460 cm^-1^ are attributed to bending modes of CH,-CH_2_, -CH_3_ that overlap with Mg-O vibration of MgO NPs given as result a decreased intensity band with respect to pure MgO. On the other hand, new bands belonging to 2ME are observed mainly at 1259 cm^-1^, 1231 cm^-1^ and 1206 cm^-1^, these peaks corresponding to the methoxy group O-CH_3_ and hydroxyl group C-OH from 2ME. Finally, in the low frequency vibration of 1179-638 cm^-1^ several bands are overlapping between 2ME and MgO-PEG which generate broader bands compared with MgO-PEG NPs. The appearance of the characteristics bands of 2ME slightly shifted when 2ME is conjugated with MgO-PEG (see Fig 3B in the range 1600-800 cm^-1^). This suggests that the interaction of 2ME and PEG may occur through hydrogen bondings of the hydroxyl groups present on the MgO-PEG surface. Furthermore, it is known that PEG is not a purely hydrophilic polymer, being able to attach hydrophobic drug as 2ME [23, 24]. For corroborate this, we perform the spectra of PEG conjugated with 2ME without MgO-NPs, this is shown in the Fig 3C. In this figure we can observe that the PEG-2ME bands are modified with respect to PEG and 2ME alone; the most striking feature of PEG-2ME spectra is that, the bands became wider than PEG bands, this may be originated from the association between OH group of PEG and hidroxyl group of 2ME by hydrogen bonding. All these characteristic and those exposed above indicate the attachment of 2ME to MgO-PEG NPs. A scheme of functionalized MgO Nps with PEG and 2ME is shown in the Fig 4.

**Fig 4.**
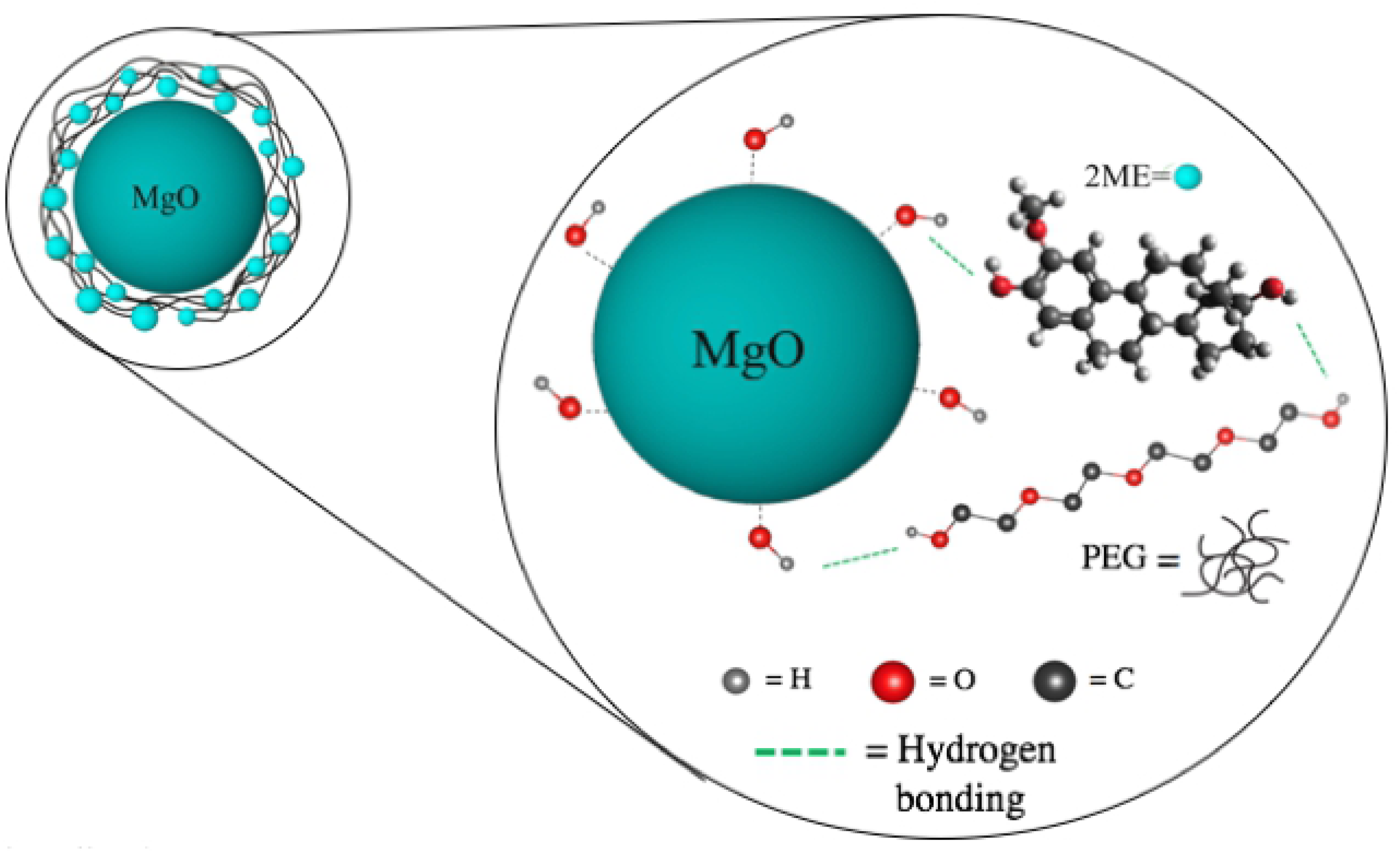
Schematic Representation. Schematic representation of MgO NPs conjugate with PEG and 2ME. The large figure shows a representation of the interaction between MgO-PEG, MgO-2ME and PEG-2ME. The Green lines represent the hydrogen bonding between OH group of PEG and 2ME molecules with OH groups of MgO NPs. The color of spheres (light gray, red and gray) represent Hydrogen (H), Oxigen (O) and Carbon (C).

### Kinetic of absorption and liberation

To evaluate the 2ME-loading capacity of the MgO-PEG NPs, we performed an extraction method in which an amount (i.e., 1 mg) of particles was dispersed in water to release the encapsulated drug, and this solution was evaporated and subsequently suspended in methanol for quantification by UPLC. The maximum absorption was reached at 3 hours of incubation with a value of 0.985 ± 0.0011 mg/ml of 2ME absorbed by each 1 mg/ml of MgO-PEG NPs that correspond to 98.51 % of absorption of total weight; this amount keep stable until 6 hours of evaluation (see Fig 5).

As the retention of the chemotherapeutic drug within the nanoparticle is fundamental for its future clinical application, we then measured the liberation of 2ME from the MgO-PEG NPs in non-biological conditions. As shown in Fig 6, we can observe that 2ME is gradually released over a period of 168 hours (7 days at pH 2, 5 and 7.35). This shows that the drug is released in a sustained manner. The mean value of release is about 33.9 %, 28.39 % and 30.16 % at pH 2, 5 and 7.35, respectively. The maximum amount released is 2.95 *µ*M that correspond to 89.27 % of the total of 2ME loaded into MgO-PEG NPs, which is reached for a pH 5 at 96 hous. For a pH 7.35, the maximum drug released was 44 % at 72 hours. These results suggest a high specificity for any potential future use of this nanoparticle since less than 1% of 2ME would be released into the circulation, and given the leaky blood vasculature that irrigates cancer cells the MgO-PEG-2ME composite should be taken up preferentially.

**Fig 5.**
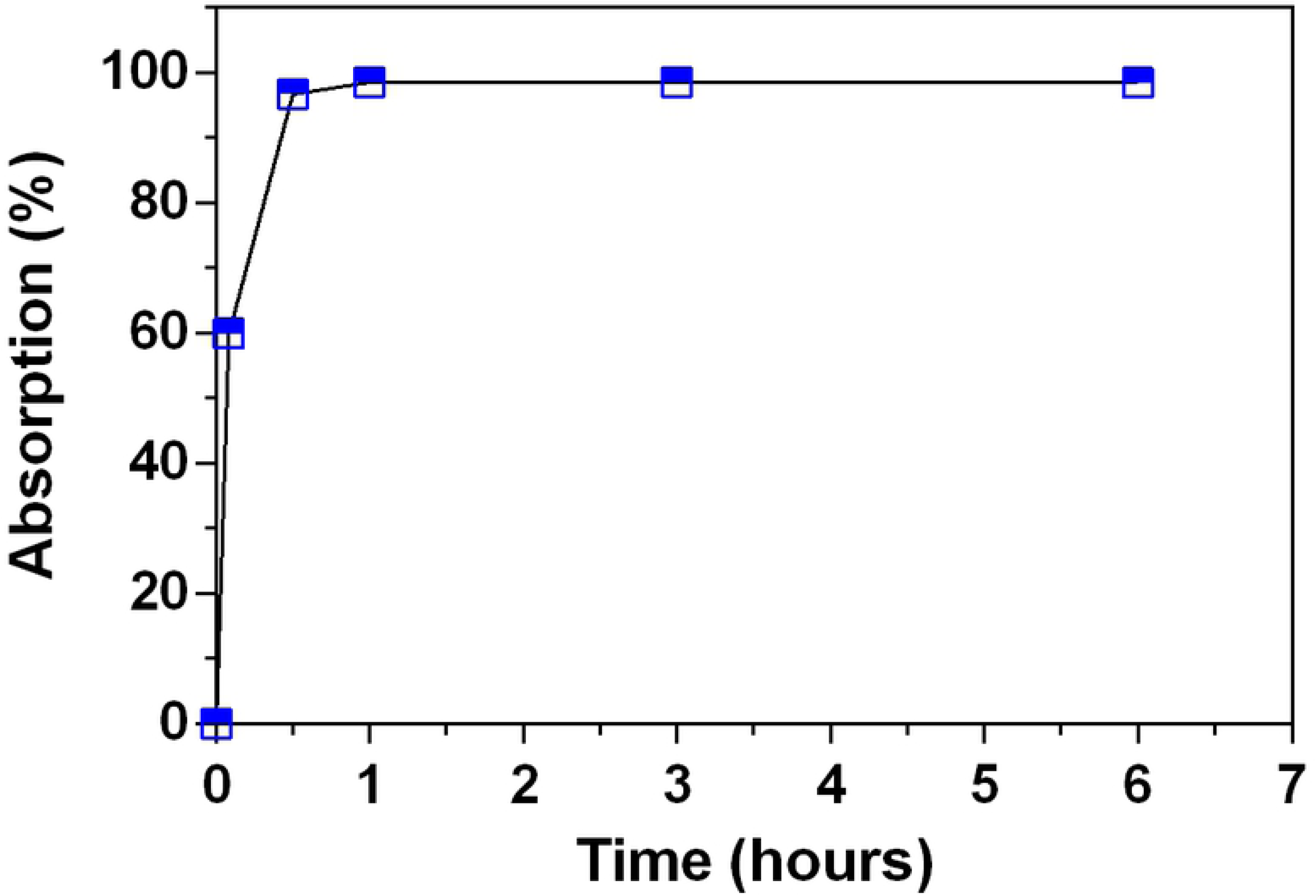
Absorption profile of 2ME by MgO-PEG NPs.

**Fig 6.**
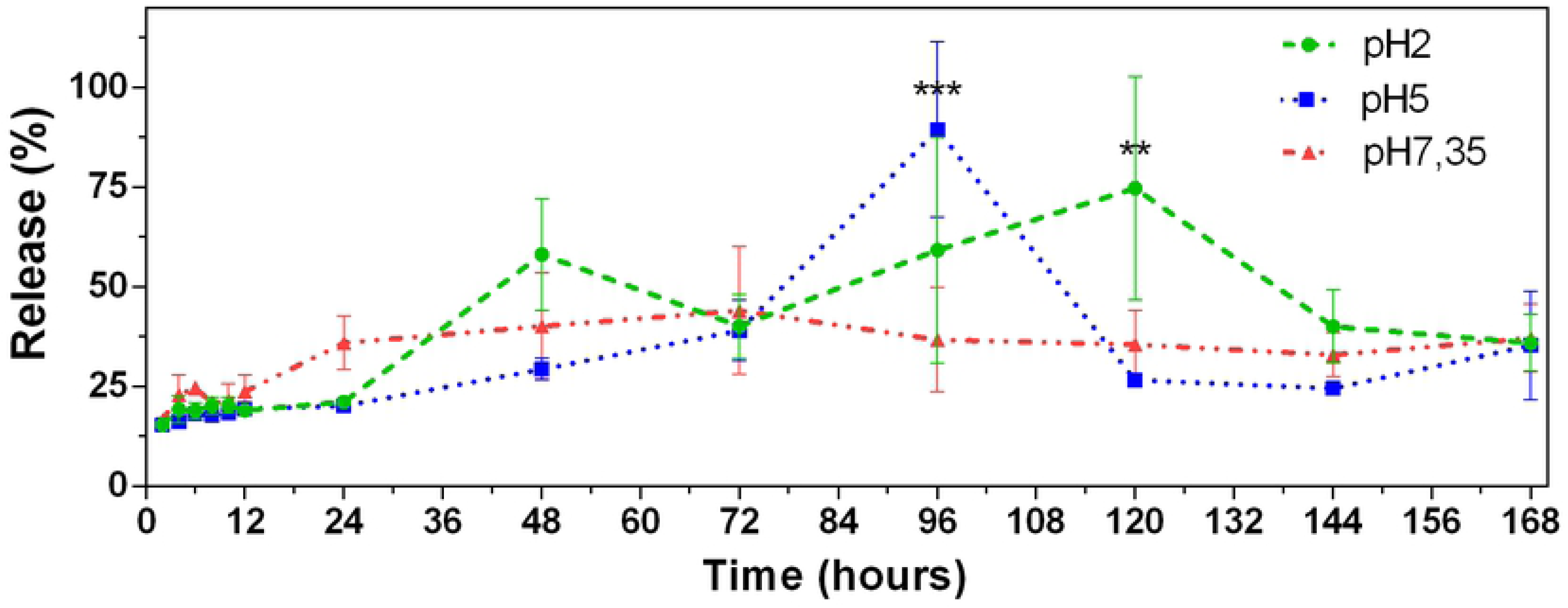
Release Profile Drug. 2ME release profile in % from functionalized MgO-PEG NPs. ** P < 0.01, *** P < 0.001

### Cell Toxicity

Viability of LnCap cells treated with MgO NPs, MgO-PEG NPs, MgO-PEG-2ME NPs at a concentration equivalent to 5 *µ*M of 2ME were determined using an in vitro viability assays (MTS). As shown in Fig 7, 2ME alone or loaded to MgO-PEG NPs induce a significant decrease in cell viability around 40% at 72 hours. These results indicate that 2ME absorbed with the MgO-PEG NPs maintains its anticancer properties suggesting that this conjugate is a promising option for therapeutic use. Interestingly, we also observed that pure MgO-NPs produce a significant decrease in the cell viability to 20% at 72 hrs, which decreases when it is coated with PEG suggesting that functionalization of MgO NPs with PEG reduces its intrinsic toxicity. This is may be explained by the fact that PEG inhibit protein absorption and/or reduce surface availability of NPs affecting their toxic activity [17, 21, 25].

**Fig 7.**
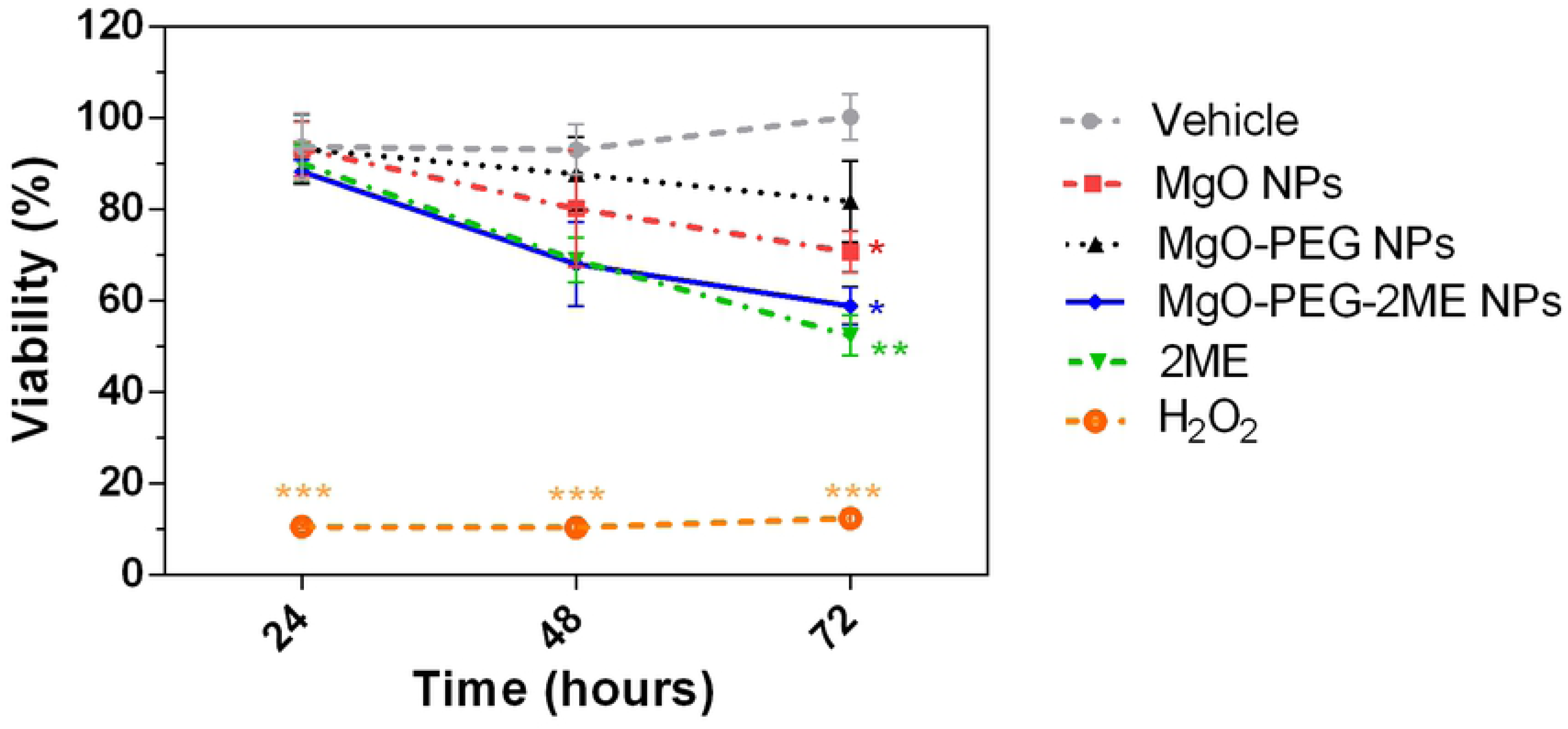
Cell Viability. Viability of LNCaP cells treated for 24, 48 or 72 hours with nanoparticles of MgO, MgO-PEG, or MgO-PEG-2ME compared with 2ME 5 *µ*M alone. Ethanol 0.01% was used as Vehicle of the nanoparticles and 2ME. As positive control of cytotoxicity we added hydrogen peroxide (H_2_O_2_) 0.08 % dissolved in 4 *µ*L of culture medium. \**P* < 0.05, ** *P* < 0.01, *** *P* < 0.001.

## Conclusion

Nanoparticles of MgO, MgO-PEG and MgO-PEG-2ME were characterized by TEM, zeta potential and FTIR spectroscopy. The modification processes attributed to the conjugation of PEG and 2ME into MgO NPs was performed step by step, where is confirmed that 2ME is attached to funtionalized MgO-PEG NPs. The 2ME absorption profile shows that a 98.51% of total weight is absorbed by MgO-PEG NPs. 2ME is released from MgO-PEG NPs constantly over a period time reaching a maximum of *µ*M at 96 hrs corresponding to 89.27 % of the 2ME total loaded into MgO-PEG NPs. In vitro viability assays (MTS) with the human prostatic adenocarcinoma cell line LNCap showed that MgO-PEG-2ME NPs has anticancer activity as similar as 2ME alone. In summary, we have developed a nanocarrier system based in MgO-PEG NPs that can load and deliver 2ME into cancer cells suggesting to this 2ME loading strategy as a promising option for use in malignant disease.The next steps of this investigation will be focused to evaluate the anticancer activity of the MgO-PEG-2ME NPs on animal models in order to assure its clinical applicability.

## Acknowledgments

This work received financial support from grants DICYT 021743OD-DAS and Basal Program for Centers of Excellence, Grant FB0807 CEDENNA, CONICYT and the DGIIP-USM. Proyecto Basal FB0821 – CONICYT. P.A. Zapata acknowledges the financial support of FONDECYT Regular N 1170226.

## Declaration of interest

The authors declare that there is no conflict of interest that could be perceived as prejudicing the impartiality of the research reported.

